# Consistency in floral volatiles impacts plant-pollinator interactions in the face of environmental change in the Eastern Himalayas

**DOI:** 10.1101/2025.02.26.640254

**Authors:** Gauri Gharpure, Joyshree Chanam, Dharmendra Lamsal, Yuvaraj Ranganathan, Sushant Potdar, Subecha Rai, Abhilasha Saikia, Adesh Rai, Dharmit Lepcha, Dharna Subba, Dorjee Tshering Lepcha, Himashri Talukdar, Jagath Vedamurthy, Megha Rai, Ongchuk Bhutia, Sapana Rai, Shrijana Chettri, Mahesh Sankaran, Shannon B. Olsson

## Abstract

Floral scents are an important mediator of many plant–pollinator interactions. However, environmental and ecological change could affect the synthesis and emission of volatile chemicals that constitute floral scents, potentially influencing pollinator perception of scents. We investigated the effects of elevation and warming on the floral scents of multiple co-occurring alpine meadow plant species along an elevational gradient in the Eastern Himalayas (3000 - 4000 masl). We also investigated pollinator responses in the same locations using 3-D printed floral models that replicated the differing volatile constituents. Finally, we assessed how pollinator preferences changed with variations in elevation, flowering season and location. We found that elevation and *in-situ* warming using Open Top Chambers (OTC) led to significant intraspecific variation in the composition of floral scents, both in terms of quality and quantity of volatiles released. Nevertheless, conspecific floral scents across elevations were more similar to each other than to other species. Cafeteria assays using artificial floral models revealed that pollinator preferences were driven by a small number of volatiles (p-cymene, 2-pentylfuran and α-pinene), but these preferences were abolished in a novel plant-pollinator community where the same floral volatiles were not detected. Our results suggest that the local odourscape established by the floral community present plays a key role in plant-pollinator interactions. This emphasizes the need to incorporate ecological impacts such as changes in community into climate change analyses to understand the long-term impacts of human activity on co-evolutionary relationships like pollination.

**Significance Statement:** Floral scents are important for most plant–pollinator interactions, and can be affected by environmental factors like elevation and temperature. Our study shows that elevation and simulated *in situ* warming change the composition of floral scents, yet scents from the same species were more similar to each other across elevations and temperatures than to other species. Further, pollinator preferences to these scents replicated in artificial models persisted in spite of variation in temperature and elevation, but not in a novel plant-pollinator community containing floral species not releasing those volatiles. This emphasizes the need to incorporate factors such as floral community composition into our understanding of the long-term impacts of climate change and human activity on co-evolutionary relationships like pollination.

## Introduction

Global environmental changes, especially climate warming, have been shown to significantly disrupt plant–pollinator interactions, often through phenological asynchrony and range shifts of both interacting partners (1, 2, 3). More recently, multiple studies have reported how climate change also impacts the volatile organic compounds (VOCs) that constitute floral scent signals (4, 5, 6, 7, 8, 9, 10, 11). With more than 80% of flowering plant species depending on insect pollinators (12), and many pollinators relying on floral scent signals to localize suitable floral resources (13, 14, 15), understanding how changes in environmental and ecological factors affect floral scents and plant–pollinator interactions is highly pertinent to discussions on climate change.

Environmental factors such as increased ambient temperature and drought are reported to affect endogenous production of certain volatiles, altering both the quantity and quality of floral scent bouquets (16, 17). Temperature also affects volatility, solubility and diffusion of VOCs so that some floral VOC emissions increase with temperature (4, 9, 10). More recent studies also show that pollutants can also modify floral VOCs, essentially impeding the olfactory cues that pollinators use to recognize flowers (18, 19). But pollinator insects can learn new modified olfactory cues to find flowers (20). Thus, while plant-pollinator interactions can be affected by multiple environmental factors, these interactions may still show a degree of robustness (3).

Alpine ecosystems such as the Himalayan region are reported to be particularly vulnerable to climatic changes such as warming as compared to other ecosystems (21, 22, 23, 24). However, there are relatively few studies on the effects of global environmental change on floral scent-mediated plant-pollinator interactions in alpine ecosystems, and even fewer in species-rich and highly vulnerable tropical alpine ecosystems such as the Himalayas. While scent signals could be affected by changing environmental factors, elevational range shifts will also have multiple and far-reaching impacts on alpine plant-pollinator interactions (24), including shifts in the plant-pollinator community (25). In this study, therefore, we investigated how plant-pollinator interactions in an alpine meadows ecosystem of the Himalayas would respond to both environmental and plant-pollinator community changes.

Mountain ecosystems, with their natural variation in community composition and temperature along elevational gradients, provide an ideal natural experimental system for climate change studies (26). Leveraging this premise, in this study we investigated the effects of elevation, temperature and flowering season on plant-pollinator interactions along an elevation gradient from 3000-4000 metres above sea level (masl) in North Sikkim in India’s Eastern Himalayan region (Figure 1). We first assessed how floral scent profiles of four species of co-occurring alpine meadow plants varied with elevation both within and across species. We also assessed the effects of changes in temperature on floral scent profiles of a fifth flower species using an *in situ* warming experiment with Open Top Chambers (OTC) that simulated increased temperatures up to 1.5-2°C. This temperature increase has been predicted to occur by the end of this present century (27), and has already been surpassed in the latest yearly global average from January 2023-January 2024 (28). In order to investigate pollinator response to varying floral scents, we conducted preliminary observations of general pollinator visitation rates to focal flowers of three of the species across elevations. We subsequently conducted behavioral assays with 3-D printed floral models releasing the differing floral VOCs emitted by these three flower species to assess the response of pollinators to floral models presented at different elevations, flowering seasons, and in a different plant-pollinator community. We discuss how changes in VOCs in these various conditions could drive pollinator preferences, as well as the implications of our results for understanding pollination in the face of environmental change in one of the most vulnerable regions on the planet.

**Figure 1.**
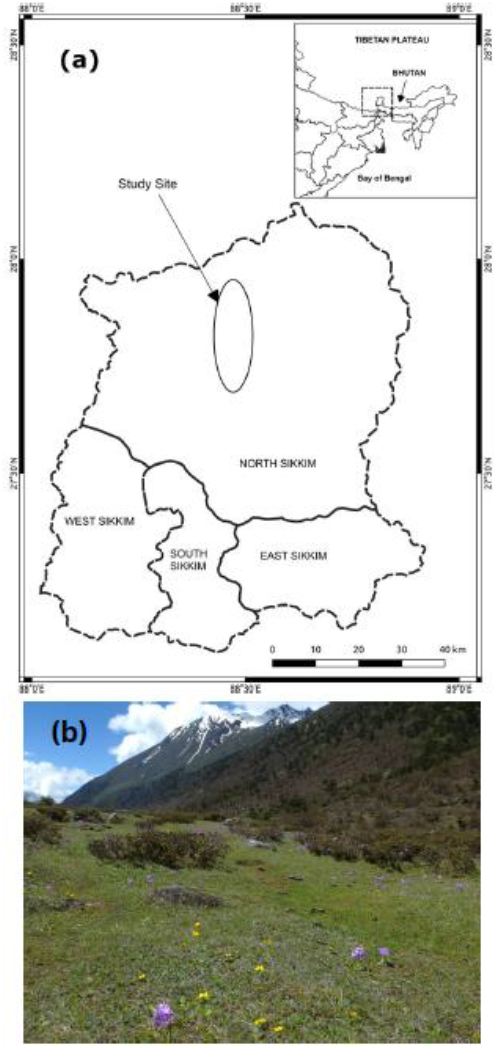
Study Location in the Eastern Himalayas. (a) Map of study site in North Sikkim (b) Photograph of one of the studied alpine meadows.

## Results

### Variation in floral VOCs across elevations

For VOC identification, volatiles were initially identified based on mass fragmentation pattern and retention time. This led to the selection of floral volatiles in *F. nubicola, A. obtusiloba*, and *L. prolifera* used for behavioral analysis (i.e. α-pinene, 2-pentylfuran, 3-carene, p-cymene, cis-3-hexenyl acetate, D-limonene, α-cedrene; Table S1.a). In further analyses, in addition to retention time and fragmentation pattern, the relative retention index (RRI) was also incorporated as a third parameter. Only those VOCs whose calculated RRI matched within 20 points of the theoretical RRI were used for the final floral VOC comparisons. This also led to the identification of a few of the initially unidentified compounds (Table S1.b). Thus, a total of 22 volatile compounds were compared across all four species across all elevations, of which 5 could not be identified and were marked as NIs (not identified) along with their respective m/z (mass/charge) in the GC-MS output (SI Appendix, Table S1.b). MDS analyses (Figure 2a-d) revealed that for each of the four flower species, floral scent profiles of individuals from the same elevation formed distinctly separate clusters, clearly showing that elevation had a significant effect on the floral VOCs within a species. These intraspecific differences in floral odours across elevations were largely driven by changes in the relative proportions of a subset of the floral VOCs detected for each species as shown in Figure 2 and listed in Table S1. When all four species were pooled together, MDS analysis (Figure 2e) revealed that floral scents of conspecifics were more similar to each other and were distinctly different from other species growing within the same community both at the same elevation and also across elevations. The VOCs pentenyl-4-cyclohexane, isocumene, longifolene, m-cymene, NI166, farnesane, NI59, 2-pentyl furan and limonene were responsible for the separate clusters across species (Figure 2e).

**Figure 2.**
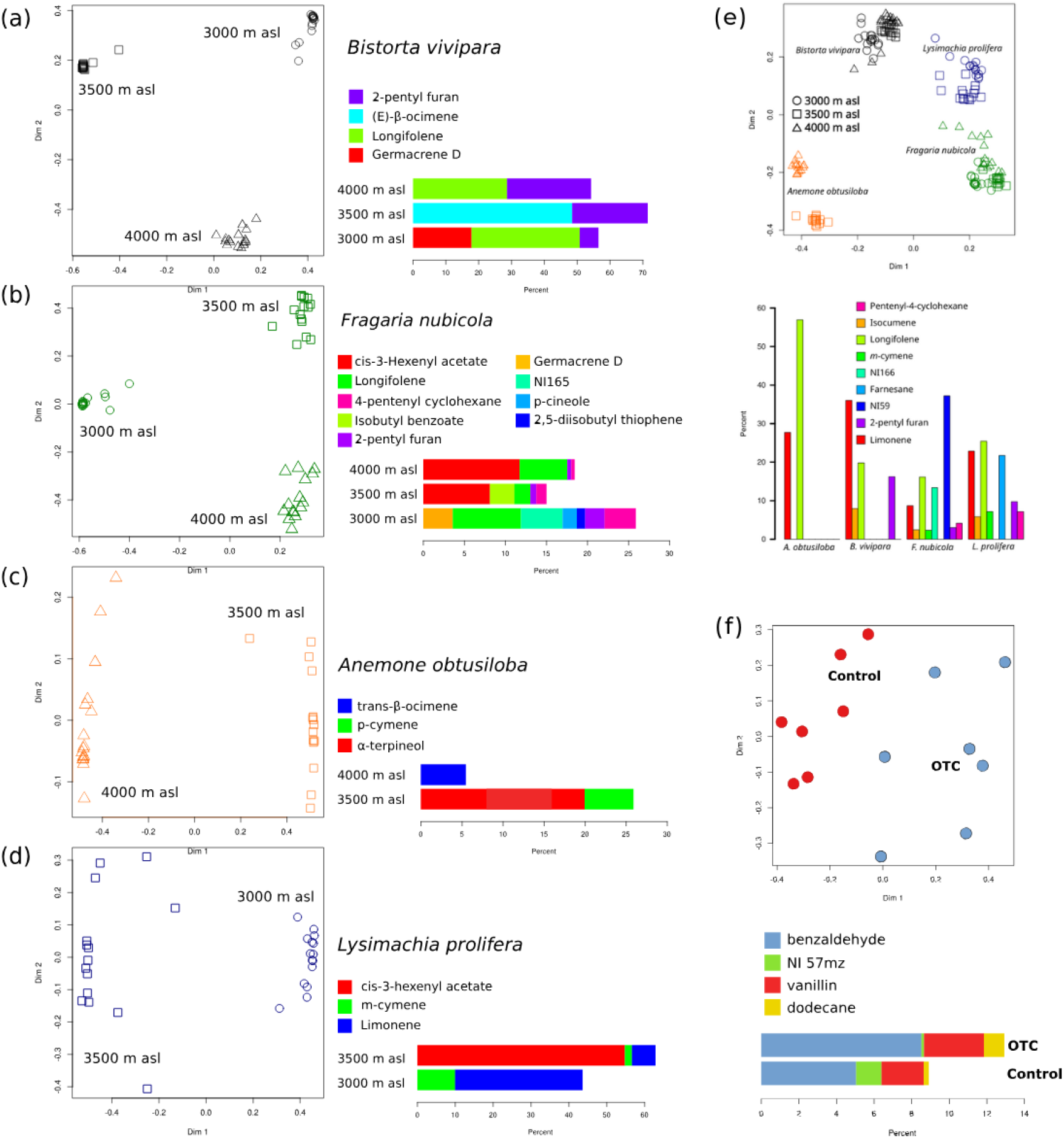
Conspecific floral VOCs form distinct clusters based on elevations. (a-d) MDS plots of four alpine flower species across different elevations; black, *Bistorta vivipara*, green, *Fragaria nubicola*; orange, *Anemone obtusiloba*; blue, *Lysimachia prolifera*. Elevations indicated by shape: circle, 3000 masl; square, 3500 masl; triangle, 4000 masl. (e) MDS plot of all four species across elevations. Shapes and colors of plots as described above. (f) MDS plot indicating floral VOC samples of *Ranunculus pulchellus* from Open Top Chambers (OTC) and control quadrats. For all plots, bar graphs depict VOCs responsible for the clustering with relative percentages in the different treatments.

### Effect of warming on floral scent profiles

Warming significantly altered the floral scent profile of *Ranunculus pulchellus*, depicted by separate clusters of floral scent samples from Warmed (OTC) and control flowers in the MDS analysis (Figure 2f). Of the total of 26 VOCs detected, four VOCs, including benzaldehyde, vanillin, dodecane and a non-identified (NI) compound NI 57m/z were responsible for the separate clustering. Warming increased the relative abundance of benzaldehyde, vanillin, and dodecane.

### Pollinator preference to floral VOCs across elevations and seasons

Our preliminary observations showed that pollinator abundance (average of total number of pollinators per quadrat) did not differ significantly across elevations (KWχ^2^ = 5.05, df = 2, p-value = 0.08). The average number of pollinator visits per flower did not vary significantly with elevation for any of the three flower species observed (*F. nubicola*: K-Wχ^2^ = 5.5324, df = 2, p-value = 0.063; *A. obtusiloba*: K-Wχ^2^ = 0.74467, df = 1, p-value = 0.39; *B. vivipara*: K-Wχ^2^ = 3.647, df = 2, p-value = 0.16). Of the three flower species, *F. nubicola* received more pollinators per flower across all elevations (number of pollinators per flower: 3000 masl: 0.486; 3500 masl: 0.667; 4000 masl: 0.098), compared to *A. obtusiloba* (3500 masl: 0.034; 4000 masl: 0.038) and *B. vivipara* (3000 masl: 0.116; 3500 masl: 0.018; 4000 masl: 0.028).

Our artificial floral models comprised of visual cues (i.e. spectral reflectance and shapes of *Anemone obtusiloba* (AO), *Fragaria nubicola* (FN) and *Lysimachia prolifera* (LP) flowers) and olfactory cues (i.e. chemicals causing separate clustering of conspecific floral VOC profiles across the three elevations viz. α-pinene, 2-pentylfuran, 3-carene, p-cymene, cis-3-hexenyl acetate, D-limonene, α-cedrene) (SI Appendix, Table S1.a). In our assays using these floral models, all models were visited by the wild pollinators across all assays (Figure 3). The models received a total of 3092 visitations across elevations, seasons and environments, with 2301 visitations in the native environment in Sikkim and 791 visitations in the novel environment in Bengaluru. Insect pollinators were identified as they visited the models and were categorised according to their respective taxonomic classes (e.g. Diptera, Lepidoptera, Hymenoptera), with flies forming the majority of visitors both in Sikkim (87.4% in May and 85% in September) and in Bengaluru (90.13%). The pollinators visited all our floral models, often extending their mouth parts to the model surface suggesting feeding behaviour, despite the lack of complete olfactory and visual cues found in real flowers, or sugar or pollen rewards.

**Figure 3.**
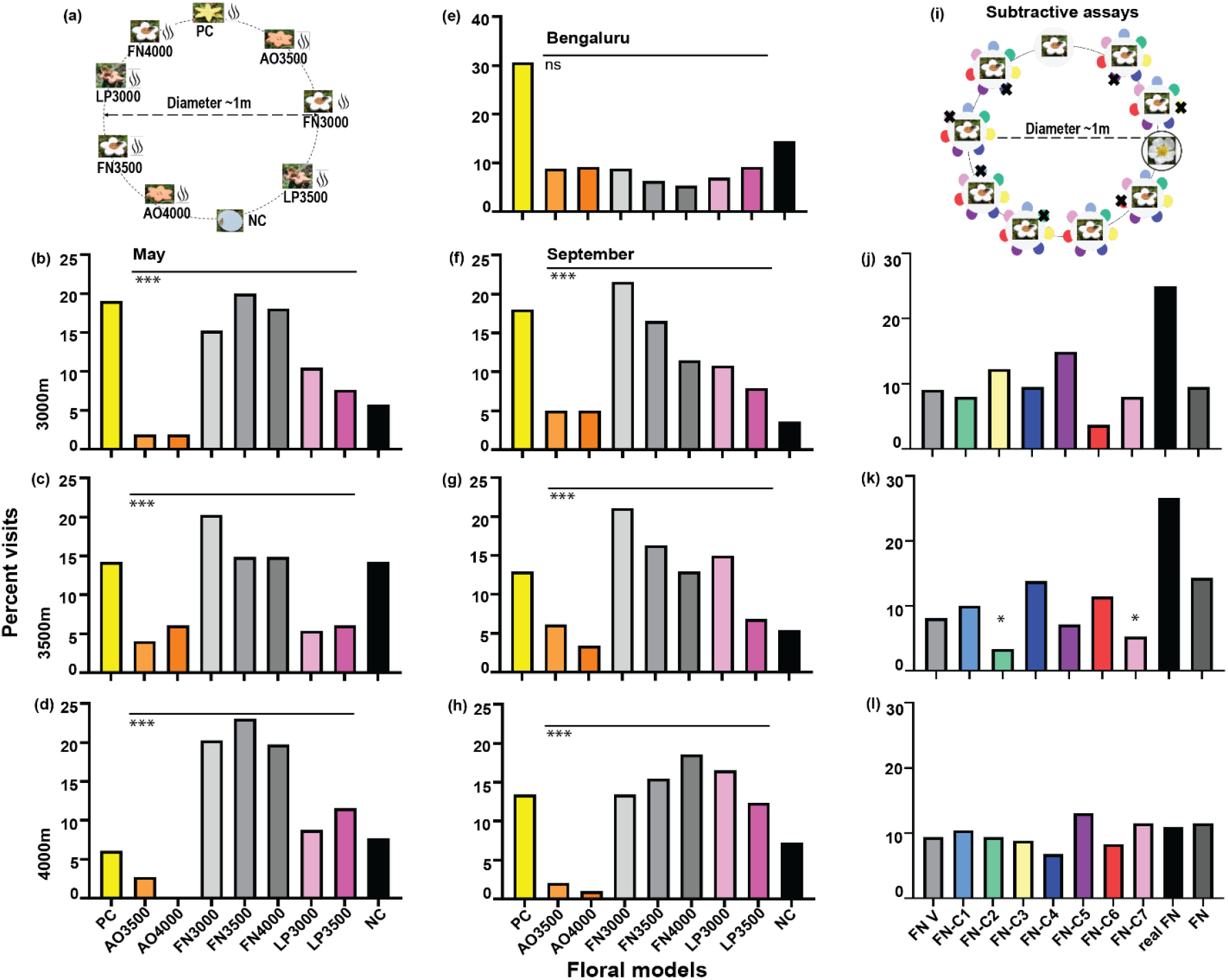
Wild pollinator visitation to artificial floral models with olfactory cues across elevations, seasons, and plant-pollinator communities. (a) Representative layout during cafeteria assays. Percent visits received in cafeteria assays in Sikkim during the regular flowering season at (b) 3000 masl, (c) 3500 masl, (d) 4000 masl. (e) Percent visits received in the novel environment in Bengaluru. Percent visits received in cafeteria assay in a different season at (f) 3000 masl, (g) 3500 masl, (h) 4000 masl (* for p < 0.05, ** for p < 0.01, *** for p < 0.001). (i) Representative layout of FN floral models during subtractive assays. Percent visits in subtractive assays in Sikkim during the regular flowering season at (j) 3000 masl, (k) 3500 masl, (l) 4000 masl. Asterisks indicate significant differences between the tested model with all chemicals and model with one chemical subtracted, based on Fischer’s exact test (* for p < 0.05, ** for p < 0.01, *** for p < 0.001). Legend: PC = Positive Control; NC = Negative Control; AO3500 and AO4000 = representative floral models of *Anemone obtusiloba* from 3500 and 4000 masl respectively; FN3000, FN3500 and FN4000 = representative floral models of *Fragaria nubicola* from 3000, 3500 and 4000 masl respectively; LP3000 and LP3500 = representative floral models of *Lysimachia prolifera* from 3000 and 3500 masl respectively; FN V = FN model with no olfactory cues; FN-C1 to FN-C7 = representative floral models of *Fragaria nubicola* with one of the seven constituent chemicals (C1 = α-pinene, C2 = 2-pentylfuran, C3 = 3-carene, C4 = p-cymene, C5 = cis-3-hexenyl acetate, C6 = D-Limonene, C7 = α-cedrene) absent; real FN = real *Fragaria nubicola* flower; FN = representative model of *Fragaria nubicola* floral model from that elevation (i.e. FN3000 at 3000 masl, FN3500 at 3500 masl and FN4000 at 4000 masl).

We noted 435 visitations to the floral models over 15 days of observation across different elevations in May (spring), with 105 visitations at 3000 masl (Figure 3 (b)), 148 visitations at 3500 masl (Figure 3 (c)) and 182 visitations at 4000 masl (Figure 3 (d)). The pollinators did not visit the models equally at the three elevations (3000 masl: χ^2^ = 33.326, p < 0.0001; 3500 masl: χ^2^ = 35.663, p < 0.0001; 4000 masl: χ^2^ = 73.198, p < 0.0001; SI Appendix, Table S5 (iv)-(vi)). These patterns of pollinator visitation also changed significantly across elevations, indicating that elevation impacted pollinator choice (3000 masl vs. 3500 masl: χ^2^ = 16.874, p < 0.0315; 3500 masl vs. 4000 masl: χ^2^ = 50.305, p < 0.0001; 3000 masl vs. 4000 masl: χ^2^ = 32.352, p < 0.0001); SI Appendix, Table S5 (xv)-(xvii).

We noted 383 visitations to the floral models over 9 days of observation in an alternate season (September, late summer) with 139 visitations at 3000 masl (Figure 3 (f)), 147 visitations at 3500 masl (Figure 3 (g)) and 97 visitations at 4000 masl (Figure 3 (h)). Again, the pollinators did not visit all models equally (3000 masl: χ^2^ = 28.221, p < 0.0001; 3500 masl: χ^2^ = 31.908, p < 0.0001; 4000 masl: χ^2^ = 25.091, p = 0.0003; SI Appendix, Table S5 (x)-(xii)). Comparing among elevations, pollinator visits at 3000 masl and 3500 masl were significantly different from visits at 4000 masl, but not between each other (3000 masl vs. 3500 masl: χ^2^ = 5.655, p = 0.6859; 3500 masl vs. 4000 masl: χ^2^ = 21.279, p = 0.0064; 3000 masl vs. 4000 masl: χ^2^ = 36.759, p < 0.0001; SI Appendix, Table S5 (xviii)-(xx)). Between the two flowering seasons, pollinator visits were similar across seasons at 4000 masl (χ^2^ = 11.039, p = 0.0872) but significantly different at 3000 masl (χ^2^ = 18.663, p = 0.0048) and 3500 masl (χ^2^ = 23.594, p = 0.0006) (SI Appendix, Table S5 (xxi)-(xxiii)). These results suggest that pollinator choices are significantly impacted by elevation and, to a lesser extent, by flowering season.

### Pollinator visitations in a unique plant-pollinator community and without olfactory cues

We noted 316 visitations to the floral models over 13 days of observation in a different environment and plant-pollinator community (i.e. subtropical Bengaluru vs. alpine Sikkim); Figure 3 (e). In this new community, pollinating insects visited the Himalayan-derived floral models almost equally (χ^2^ = 6.628, p = 0.3566); SI Appendix, Table S5 (xiii), (xiv). On sampling floral VOCs from the nearby fully-bloomed flowers within 5 metres of the assay following the protocol described in methods, we did not detect any of seven constituent chemicals in any flowers in the vicinity (SI Appendix, Table S6). Interestingly, pollinators did not visit the floral models without olfactory cues equally (χ^2^ = 17.958, p = 0.0063) (SI Appendix, Figure S3, Table S5 (xiii)). This is in contrast to Sikkim, where pollinators visited the floral models without olfactory cues unequally at 3000 masl (χ^2^ = 54.379, p < 0.0001) and 4000 masl: χ^2^ = 61.348, p < 0.0001), but not at 3500 masl (χ^2^ = 7.4000, p = 0.2854) (SI Appendix, Figure S3, Table S5 (i)-(iii)). In Sikkim, the pollinator visitation patterns to floral models without olfactory cues were similar to those shown towards floral models with olfactory cues at 3000 masl (χ^2^ = 9.052, p = 0.3379) and 4000 masl (χ^2^ = 15.068, p = 0.0578), and not at 3500 masl (χ^2^ = 24.562, p = 0.0018); SI Appendix, Figure S3, Table S5 (xxiv)-(xxvi). These results suggest that pollinators can choose among models without olfactory cues in multiple environments. However, these pollinators also choose among floral models with olfactory cues only in the presence of the floral community and volatiles in Sikkim, and not in their absence in the novel plant-pollinator community in Bengaluru.

### Effect of specific floral VOCs on pollinator visitation

We presented wild pollinators with FN floral models containing olfactory cues comprising of all but one constituent chemicals of the blend of that elevation (e.g. FN3000 blend at 3000 masl). In these subtractive assays, all floral models at all elevations were visited by wild pollinators, with 188 visitations at 3000 masl (Figure 3 (j)), 210 visitations at 3500 masl (Figure 3 (k)) and 192 visitations at 4000 masl (Figure 3 (l)). At 3000 masl and 4000 masl, all floral models were visited similarly to the positive controls. However, at 3500 masl, absence of 2-pentylfuran and α-cedrene from the blend caused a significant reduction in the number of visitations (Fischer’s exact test, p = 0.0068 and p = 0.0408 respectively). This indicates that for FN models at 3500 masl, 2-pentylfuran and α-cedrene are important to receive visitations.

Our GLM results (SI Appendix, Table S7) showed that α-pinene, 2-pentylfuran and p-cymene, were significant in impacting visitations to all seven AO, FN and LP models across elevations. Regression tree analyses (SI Appendix, Figure S4) showed similar results to GLM for the seven models. The presence of p-cymene was the most important factor for pollinator visitation across elevations, while α-pinene and 2-pentylfuran determined visitations at certain elevations. The next important factor was found to be the identity of the model, with FN models consistently receiving most visitations as shown for real flowers in our preliminary assays. Thus, the presence of the 3 chemicals in sufficient quantities and the identity of the floral model were found to contribute significantly in receiving pollinator visitations to AO, FN and LP models at all three elevations.

## Discussion

Our results across various environments demonstrate that ecological factors such as plant-pollinator community composition impact chemically-mediated plant-pollinator interactions more than factors such as elevation and temperature. Floral odour profiles were more similar to one another than to other species found within the same location, both within and across elevations and temperatures. Further, wild pollinator preferences for these volatiles were retained across elevation and flowering season, but not in a novel plant-pollinator community containing floral species not releasing those volatiles.

This is not to say that factors like elevation and temperature do not impact plant-pollinator interactions. Indeed, intraspecific variation in the floral odour profiles of our four tested montane meadow species showed differences across elevations in both the quantity and quality of volatile constituents released in the floral headspace. These differences were largely driven by changes in mono- and sesquiterpenes, although we detected no consistent patterns with respect to elevation. Our experiment with *R. pulchellus* using OTCs also confirms that even in alpine ecosystems where the ambient temperature through the growing season is much lower (11.3°C ±4.4 SD), increased ambient temperatures of as little as 2°C can significantly alter floral odour composition. Increased temperature has been known to increase volatility and emission rate of floral VOCs (4, 9, 10, 11), especially terpenes within the temperature range of 25–35°C, when temperature of just the flower headspace was increased by 5 °C for 10 minutes, in Mediterranean plants (6). Nevertheless, our results indicate that inter-specific differences in floral odours override intraspecies variation. Floral odour profiles of a given species across elevations were more similar to one another than to other species both within and across elevations.

When replicating the differences in floral odour profile across elevations using artificial models, pollinators were able to distinguish and choose particular odour profiles, indicating that these odour profiles are distinguishable by the resident pollinators. Chemicals such as p-cymene and 2-pentylfuran were found to be important for this segregation. Moreover, *Fragaria nubicola* flowers and the models replicating this species always received most visitations, indicating that the identity of the species is important. Nevertheless, these preferences were completely lost in a novel plant-pollinator community where the floral species present do not release these volatiles in detectable quantities. Therefore, the local chemical milieu of the floral community has a significant impact on the preferences of pollinators.

Such a phenomenon supports a chemical component to floral constancy (38), wherein a pollinator, once having found and associated a certain flower odour signal or morphotype with rewards, will visit flowers having similar signals in the midst of intraspecies variation in VOC profiles. This could provide the pollinators with a mechanism to compare floral odours of different species with each other in the local community to identify target flowers in a manner analogous to the comparison of wavelengths for colour constancy. Such a phenomenon could also explain why pollinators in our study exhibited preferences to our artificial models at different elevations and in flowering seasons, but not in a new plant-pollinator community where those floral volatiles were not detected in the species present. These results are also supported by other studies where context was crucial for pollinators to distinguish among similar odour cues (39). This suggests that the local odourscape established by the floral species present plays a key role in plant-pollinator interactions (40).

Tropical mountain ecosystems such as the Himalayas are biodiversity hotspots expected to suffer greater impacts of increased future temperatures when compared to lowland as well as temperate ecosystems. With increased warming, life forms from lower elevations are expected to migrate upwards, while high altitude alpine communities potentially face threats of extinction due to increased warming (41). Our results indicate that floral odours will likely change in a warmer future climate, specifically in the composition and relative proportions of the VOCs within a species. However, our results also suggest that changes in floral community composition and the local odourscape created by that community could have an even greater immediate impact on plant-pollinator interactions than environmental changes.

Beyond climate change, the Himalayan region, like most ecosystems in the Global South, is experiencing profound changes due to land development, environmental degradation, biodiversity loss, agricultural displacement, landscaping, and invasive species. Effects of such large ecological changes on plant-pollinator interactions have not been well explored (1). Our results suggest that such changes in floral community composition can have potentially disastrous consequences like loss of floral preference of pollinators. Fortunately, such changes can be addressed through efforts in ecosystem restoration, conservation, and wildlife corridors, among many other methods. In the lens of the Anthropocene, it is important to look beyond our carbon tunnel vision to address the many other ecological impacts humans are having on this planet (42), which could be having much more immediate and long-lasting effects on co-evolutionary relationships like pollination.

## Materials and Methods

### Study site

Our study was conducted in the alpine meadows of North Sikkim district, Sikkim state, in the Eastern Himalayas of India (Figure 1a), a global biodiversity hotspot (23). This region lies within the Kanchenjunga Biosphere Reserve, listed in UNESCO’s World Network of Biosphere Reserves. The alpine meadows (Figure 1b) of this region extend from 3000 masl to around 4500 masl, above which vegetation becomes sparse, and eventually barren. Our study was conducted at three different elevations: 3000, 3500 and 4000 masl. Average summer temperatures during our study period were 12.6 °C (±3.4 SD) at 3000 masl, 11.3 °C (±4.4 SD) at 3500 masl and 10.4 °C (± 4.1 SD) at 4000 masl.

### Floral VOC Collection

We sampled floral VOCs from four plant species distributed across our study range during June-August 2015 and June-August 2016. *Bistorta vivipara* (Polygonaceae) and *Fragaria nubicola* (Rosaceae) were found at all the three elevations (3000, 3500 and 4000 masl), while *Anemone obtusiloba* (Ranunculaceae) was found at 3500 and 4000 masl, and *Lysimachia prolifera* (Primulaceae) was found at 3000 and 3500 masl. We chose freshly opened flowers of similar phenological stage, determined visually, to collect VOC samples. For each species, we sampled 15 replicate flowers at each elevation, except for *F. nubicola* and *L. prolifera* at 3500 masl for which we sampled 16 replicate flowers each. Floral VOCs were collected *in situ* following a protocol described in (30), which was itself modified from (29) (Supplementary Material S1).

### *In situ* warming experiment

To investigate the effect of warming on floral scents, we leveraged an ongoing alpine grassland warming experiment established at our study site at 3500 masl in 2011. These experimental plots were warmed using passive Open Top Chambers (OTCs) that are often used to study global warming effects (31, 32, 33, 34). Our OTCs were constructed as described in SI Appendix, Materials and Methods. Control plots (1 m x 1 m) were established adjacent to the OTCs. OTCs were set up every year at the end of snow-melt (April) before underground rhizomes gave rise to new shoots, and dismantled at the end of the growing season in October. We used *Ranunculus pulchellus* (Ranunculaceae) at 3500 masl as the model as it was the only species for which we had sufficient number of flower replicates naturally growing in both OTCs and control plots, i.e., one flower each from seven different plants inside seven OTCs, and one flower each from seven different plants in control plots adjacent to the OTCs. The seven OTC and control pairs were spread across three plots at the same elevation, each plot having four OTCs and four control quadrats.

### Pollinator response to varying floral scents

For the purposes of this study, we classified all flower-visiting insects as pollinators. We first conducted preliminary observations to estimate pollinator abundance across elevations as well as pollinator visitation rates on focal flowers for the different flower species across elevations (SI Appendix, Materials and Methods). We then performed behavioral assays to determine how wild pollinators responded to changing floral scents using 3-D printed flower models.

Artificial floral models were constructed using a previously standardized method in our lab group (35) (SI Appendix, Materials and Methods). A total of seven floral models with specific VOC blends were standardised from three flower species across three elevations, including *Anemone obtusiloba* from 3500 and 4000 masl, *Fragaria nubicola* from 3000, 3500 and 4000 masl, and *Lysimachia prolifera* from 3000 and 3500 masl, referred to as AO3500, AO400, FN3000, FN3500, FN4000, LP3000 AND LP3500 respectively (SI Appendix, Table S5).

We used the aforementioned seven floral models along with a positive control (PC-floral object previously identified as a universal generalist pollinator attractant in the same locations (30)) and a negative control (NC - an odorless grey disc) in the field assays to determine pollinator response. In each assay, floral models were arranged in a circle of 1 metre diameter (Figure 3 (a)), which has previously been shown to prevent mixing of VOCs from adjacent floral models (30). A 5μl standardized blend was added to fresh PCR tubes in each floral model just before starting the observations. The order of models in the pair was randomised every day, with the positive control and negative control always diametrically opposite to one another (see 30).

Pollinator visitations were characterised both by insects landing on the surface or showing directed flight within 15 cm of the model surface (30, 35). Observations were made over two hours during peak pollinator activity time determined during pilot experiments. Equivalent models without floral odours were also used to compare the effect of odour on pollinator behaviour. The assays were performed in pairs with one observer monitoring per assay, and observers exchanged the assay after one hour to account for observer bias. To assess the impact of elevation, cafeteria assays were performed at 3000, 3500 and 4000 masl over 15 days in May 2022. Cafeteria assays were also repeated at the same elevations over 9 days in September 2021, when other flower species were predominant, yet a few species, such as *Anemone* sp., *Fragaria nubicola*, and *Taraxacum* sp. were still present in some locations. Preliminary observations found several similar pollinator species present in both seasons, such as hoverflies, e.g. *Eristalis tenax*, other *Eristalis* sp., *Episyrphus balteatus, Scaeva* sp., tabanid flies, muscid flies and blowflies. To assess pollinator response to the same 9 artificial floral models in a novel plant-pollinator community, we also performed the behavioural assays in Bengaluru over 13 days in September-October 2020 where the aforementioned floral species have not been known to occur, and the pollinator community is also different. A total of 20 observers performed these cafeteria assays over 37 days in Sikkim and Bengaluru.

To investigate the effect of specific VOCs on visitations, we also performed cafeteria assays using a subtractive approach. The *Fragaria nubicola* (FN) models received a majority of the visitations at all three elevations and were used for this subtractive assay. This assay involved removing one volatile constituent from each of seven FN models (Figure 3 (i)). In this assay, the positive control was the model with olfactory stimuli comprising the complete blend of that elevation, while the negative control was the FN model with no olfactory stimuli (i.e., a model with only the visual stimuli). A real *Fragaria nubicola* flower was also included in the array for comparison. These assays were performed at each elevation by 6 observers over 8 days in May 2022.

### Data analysis

Volatile organic compounds were identified by comparison with synthetic standards and/or through a comparison of their retention index and the NIST MS library (National Institute of Standards and Technology) after the peaks were deconvoluted using the AMDIS software (Automated Mass Spectral Deconvolution and Identification System). We used octamethylcyclotetrasiloxane (RRI: 991; Base peak: 281 m/z) as the internal standard (29, 30, 36) and all VOCs were represented as relative ratios to this compound. Of the total number of compounds identified for each species at a given elevation, only those compounds present in more than 75% of the samples were analyzed further. Multi-dimensional scaling (MDS) analysis was conducted using compound identities and relative peak areas to visualise the similarity/dissimilarity of data points. Each data point represented the combination of VOCs in multidimensional space. Chi-square tests of independence were used to examine if all models were visited equally in the cafeteria assays as well as between elevations and seasons, where a statistically significant chi-square value indicates pollinator choice. Fischer’s exact test was used to determine which model received significantly different visitations against the positive and negative controls in the subtractive assays. Generalised Linear Models (GLM) were performed to determine which chemical drives visitations to the different floral models across elevations, with the number of insect visitations to the floral model as the dependent variable and the amount of each chemical (in microlitres) as the independent variable with a Poisson distribution and log link function. Regression tree analysis was also employed with the amounts of all chemicals as covariates and number of insect visitations to the floral model as the response variable. We pruned the terminal nodes to ensure minimum mean square error (MSE) and complexity parameter (CP) values resulting in the most parsimonious tree. All analyses were carried out using R software (37) and GraphPad Prism software 8, GraphPad 303 Software. Inc, California, USA.

## Acknowledgments

The authors thank Chengappa S.K., Siddharth Machado, Nandita Nataraj, Nawang Hissay Lachenpa, Bhagta Bahadur Subba and Sanu Bhai for assistance in field work, Chintan Sheth for preparing the map of study site, V.S. Pragadheesh and Srinivas Rao for help with the sample analysis, Anal Kumar, Harshavardhan, Hinal Kharva, Mayuresh Gangal, Mihir Joshi, Mukta Mande, Praveen Prakash and Sulu Mohan for helping in data collection in the Bengaluru experiments in spite of the Covid-19 restrictions in 2020. The authors would like to acknowledge Aditi Mishra, Deepa Rajan, Geetha GT, Hinal Kharva, Mayuresh Gangal, Nachiket Kelkar, Nandita Nataraj, Pavan Kaushik, Priyadarshini Gurung and Shweta Basnett for insightful discussions on the manuscript. JC thanks Guido Caniglia, Isabella Sarto-Jackson and Gerd B. Müller for care and support during the final stage of the manuscript. We also thank the people of Sikkim, especially Nawang Pinsho Lachenpa for logistics support and hospitality during the field experiments in Sikkim as well as the Sikkim Forest Department, Pipon and Lachen Dzumsa, Sikkim Police Department, Indo-Tibetan Border Police, Indian Army, Home Department for permits and support.

