## Supplementary Information for "Consistency in floral volatiles impacts plant-pollinator interactions in the face of environmental change in the Eastern Himalayas"

**This PDF file includes:**

- Supporting text
- Figures S1 to S4
- Tables S1 to S6
- Legends for Datasets S1 to S2
- SI References

**Other supporting materials for this manuscript include the following:**

- Datasets S1 to S2

### SI Appendix

### SI Materials and Methods

**Floral VOC Collection:**

Floral Volatile Organic Compounds (VOCs) were collected *in situ* following a protocol modified from (1), and described in (2). Briefly, Polydimethylsiloxane (PDMS) tubes (Carl Roth Rotilabo, silicone tube; length 5mm, 1.5 mm inner diameter and 3.5mm outer diameter), were used for passive volatile collection. Tubes were first conditioned by placing them in 1:1 acetonitrile/methanol for 3 h, drying under nitrogen, and then heating from 40 °C to 260 °C at 10 °C/min, and held at 260 °C for 160 min under 4 bar nitrogen in a Tube Conditioner (Gerstel). These PDMS tubes were then cooled to 25 °C. Conditioned PDMS tubes were placed in amber vials and flushed with nitrogen, and capped and stored at −20 °C until use. To collect floral VOCs, for each flower, two conditioned PDMS tubes strung on a bent steel wire were suspended inside a transparent plastic cup. The plastic cup was covered with a transparent UV filtering sheet to protect floral VOCs from direct sunlight and wind. The steel wires and cups were pre-cleaned with 100% ethanol. The PDMS tubes were suspended approximately 1cm above the flower positioned inside the cup. Floral volatiles were collected by passive adsorption on the PDMS tubes for 4 hours, following which the tubes were removed and placed in amber vials and sealed with Teflon tape and stored at 4 °C until analysis. To account for possible contamination during sampling, blank tubes subjected to the same transportation and storage regimes were analyzed along with the sampling tubes.

**Floral VOC Analysis:**

The VOC analysis was performed by following the protocol established previously in (2). Floral VOCs were analysed by Gas Chromatography-Mass Spectrometry (GC-MS) connected to a Gerstel Thermal Desorption Unit (TDU). Desorption of volatiles employed a Gerstel TDU in a splitless mode and a Cooled Injection System (CIS 4) controlled by Gerstel Modular Analytical Systems Controller C506 and software Gerstel Maestro 1. PDMS tubes were inserted into the TDU at 30 °C using a Gerstel Multi-Purpose Sampler. After a 1min delay at 30 °C, the temperature of the TDU was increased to 200 °C at the rate of 100 °C/min, and held at 200 °C for 10 min. The desorbed volatiles were transferred at 210 °C and trapped in a silanized glass wool liner of the CIS at −50 °C using liquid nitrogen. After a 0.20 min of equilibration time, the CIS was ramped to 220 °C at a rate of 12 °C/s and held for 5 min. Volatiles were separated and identified in solvent vent mode with a purge flow to split vent of 30 mL/min at 1.5 min and vent flow of 70 mL/ min, and vent pressure at 7.07 psi for 0.01 min using an Agilent 7890B gas chromatograph coupled with a 5977A MSD mass spectrometer using an HP-5 MS column (30 m × 0.25 mm i.d., 0.25 µm film thickness) with helium carrier gas at 1 mL/min. The column oven was kept at 40 °C for 1 min, increased to 180 °C at 5 °C/min with a 5-min hold, and finally increased to 270 °C at 25 °C/min. Compounds were identified by MS in electron impact mode with ionization energy of 70 eV, a transfer temperature of 250 °C, and source and quadrupole temperatures of 230 °C and 150 °C, respectively. Different volatile analytes were identified by mass spectrometry (MS) using Agilent MassHunter Workstation software B.07.02.1938. MassHunter Qualitative Analysis, version B.07.00 was used to verify the component identity by matching the obtained mass spectral data and spectra in National Institute of Standards and Technology library, as well as injection of synthetic standards.

***In situ* warming experiment:**

Our Open Topped Chambers (OTCs) were four-sided enclosures made of acrylic sheets, and open at the top to allow gas flux so that there was no difference in air composition within and outside the OTCs. The OTC walls were 0.5 metre tall with an inward inclination of 60°, such that a quadrat of 1 metre x 1 metre fitted directly below the open top, and increased the inside temperatures by ~1.5-2 °C above ambient, which is similar to the projected increase in global average temperature by the end of the present century (IPCC 2014, 2018).

**Pollinator abundance and flower visitation rates:**
The visitation rate of pollinators to a flower was used as a measure of attraction to the flower. Pollinator counts were performed using a 1m x 1m portable quadrat frame. The quadrat frame was randomly placed for each sampling point. The total numbers of pollinators in the quadrat, as well the number of pollinators that visited a focal flower within the same quadrat within a span of 2 minutes were noted to estimate pollinator abundance and pollinator visitation rate respectively. For our analysis, we used only those quadrat samples that had only one flower species to avoid confounding observations, as we did not know whether different flower species were equally attractive to pollinators or not. Pollinator visitation data was collected for three of the four flower species, viz., *Anemone obtusiloba* (Ranunculaceae), *Fragaria nubicola* (Rosaceae) *and Bistorta vivipara* (Polygonaceae) as they occurred within our random quadrats. We sampled a total of 109 quadrats across the three elevations (3000 masl: 37 quadrats; 3500 masl: 28 quadrats; 4000 masl: 44 quadrats).

**3-D floral models:**
Artificial floral models were constructed using a previously standardized method in our lab group (3). Briefly, visual cues like floral dimensions and spectral reflectance of real flowers were available from a previous study conducted in the same landscape (2). Floral models were designed in OPENSCAD software and 3-D printed using polylactic acid (PLA) and painted with a standardized paint blend (Table S2) to have the spectral reflectance as close to that measured from the real flowers (Fig. S1, S2). The olfactory cues comprised of the VOCs which exhibited intraspecific variation across elevations. A total of 7 chemicals (alpha-pinene, 2-pentylfuran, 3-carene, cis-3-hexenyl acetate, p-cymene, D-limonene and alpha-cedrene; Table S3) were used for standardizing the blends by matching the relative ratios of the percent emission with that obtained from PDMS sampling of floral VOCs collected from flowers at each elevation (Table S4). The standardized blends were added to a PCR tube placed inside the painted floral model. The set-up was equilibrated following previously established protocol in (2) (described in detail below). A total of seven floral models with specific VOC blends were standardized from three flower species across three elevations, including *A. obtusiloba* from 3500 and 4000 masl, *F. nubicola* from 3000, 3500 and 4000 masl, and *L. prolifera* from 3000 and 3500 masl, referred to as AO3500, AO400, FN3000, FN3500, FN4000, LP3000 AND LP3500 respectively (Table S5).

**Standardization of floral blends:**
5µl of the blend was added to a PCR tube placed inside the painted floral model. The set-up was kept for equilibration for 30 minutes by covering the beaker with aluminium foil. A 100 µm PDMS+DVB+CAB SPME (solid-phase microextraction) fibre (model no:548575-U, SUPELCO, USA) was exposed to the models for 10 minutes. The exposed fibre was immediately injected into a GC-MS column in Agilent 7890B gas chromatograph coupled with a 5977A MSD mass spectrometer using an HP-5 MS column (30 m × 0.25 mm i.d., 0.25 µm film thickness) with helium carrier gas at 1 mL/min. The temperature was initially kept constant at 40°C for 1 min, increased from 40° to 100°C at the rate of 2°C per min, then ramped from 100° to 200°C at 10°C per min, and a final ramp from 200° to 270°C at 30°C per min. The temperature was then kept constant at 270°C for 5 min. The total run time was 47.6 min. The percent emission of each volatile analyte was quantified by calculating the area under the peaks of relevant compounds.

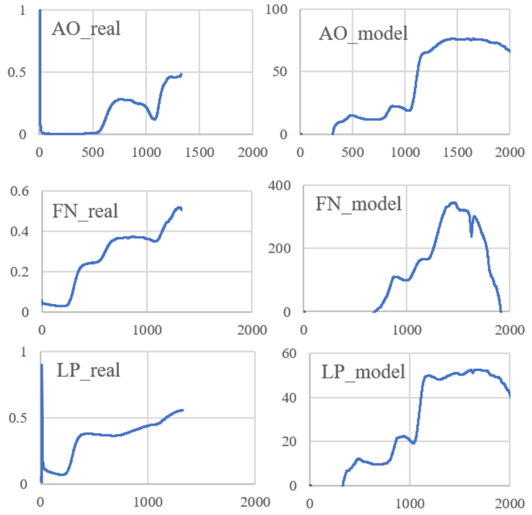


**Fig. S1.** Spectral reflectance of floral models, where AO is *Anemone obtusiloba*, FN is *Fragaria nubicola* and LP is *Lysimachia prolifera*.

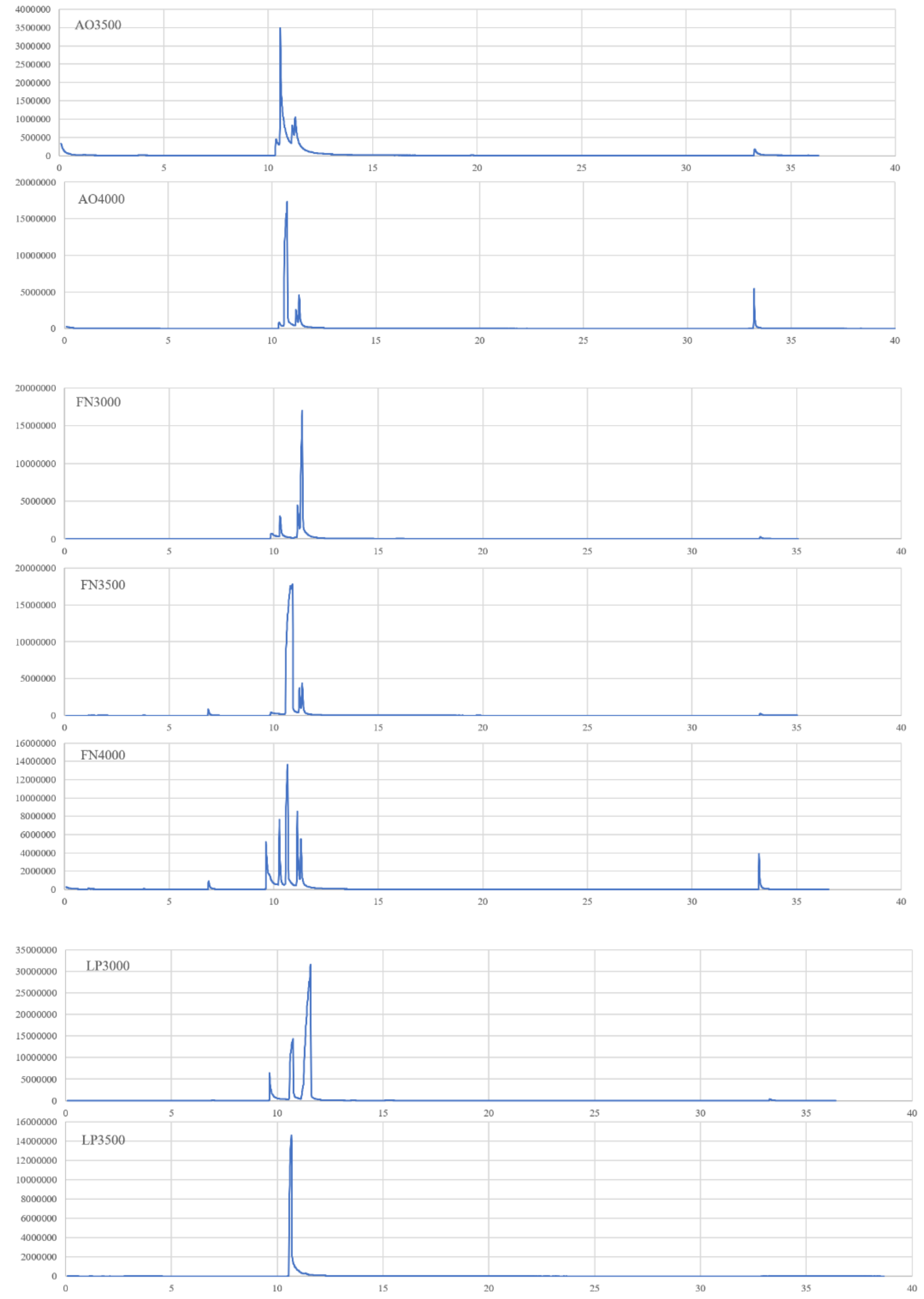


**Fig. S2.** Gas Chromatography-Mass Spectrometry (GC-MS) traces of the blends. Legend: AO3500 and AO4000 = representative floral models of *Anemone obtusiloba* from 3500 and 4000 masl respectively; FN3000, FN3500 and FN4000 = representative floral models of *Fragaria nubicola* from 3000, 3500 and 4000 masl respectively; LP3000 and LP3500 = representative floral models of *Lysimachia prolifera* from 3000 and 3500 masl respectively.

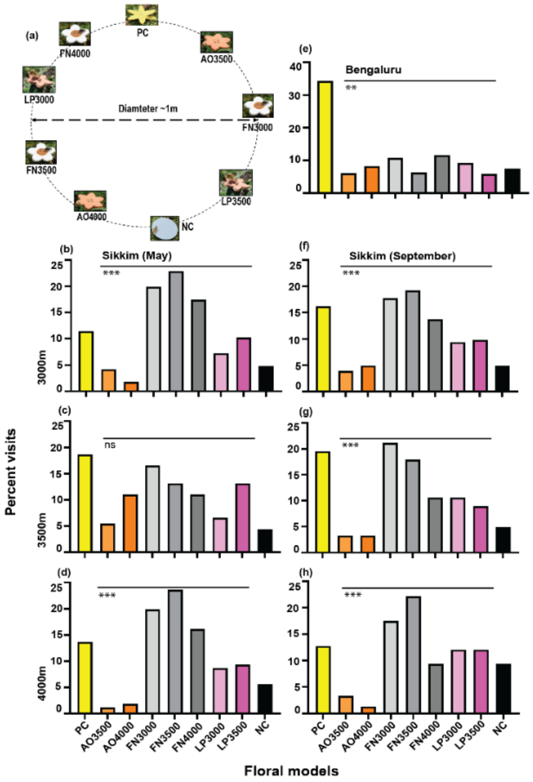


**Figure. S3.** Wild pollinator visitation to artificial floral models without olfactory cues across elevations, seasons, and plant-pollinator communities. (a) Representative layout during cafeteria assays. Percent visits received in cafeteria assays in Sikkim during the regular flowering season (May) at (b) 3000 masl, (c) 3500 masl, (d) 4000 masl. (e) Percent visits received in Bengaluru. Percent visits received in cafeteria assay in Sikkim in a different season at (f) 3000 masl, (g) 3500 masl, (h) 4000 masl (* for $p < 0.05$, ** for $p < 0.01$, *** for $p < 0.001$. Stars indicate significant differences based on $\chi2$-test of independence (* for $p < 0.05$, ** for $p < 0.01$, *** for $p < 0.001$). Legend: PC = Positive Control; NC = Negative Control; AO3500 and AO4000 = representative floral models of *Anemone obtusiloba*; FN3000, FN3500 and FN4000 = representative floral models of *Fragaria nubicola*; LP3000 and LP3500 = representative floral models of *Lysimachia prolifera*.

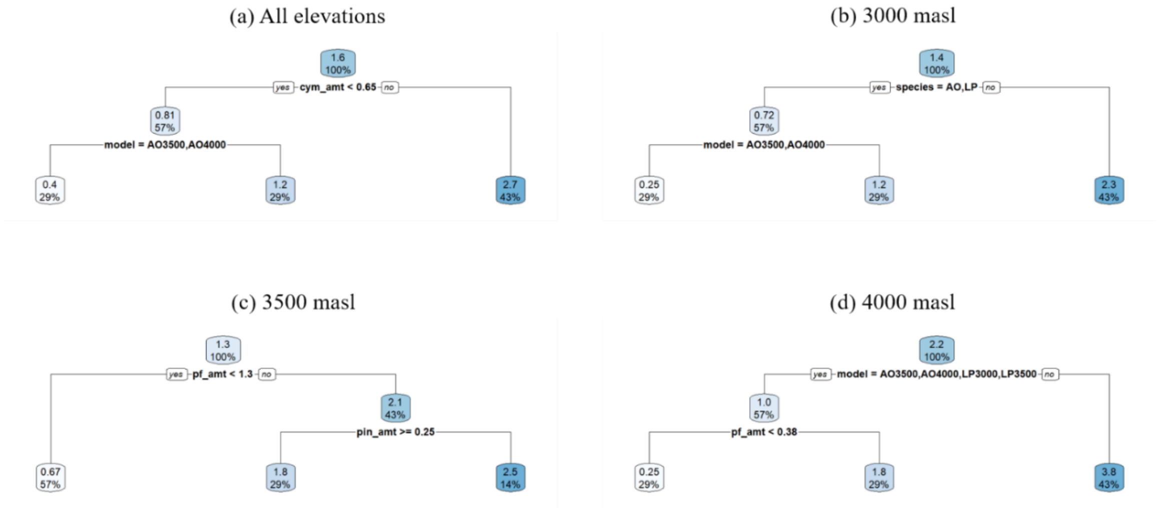


**Fig. S4.** Regression trees showing the factors and their extent of contribution to wild pollinator visitations to artificial floral models at (a) all elevations, (b) 3000 masl, (c) 3500 masl, and (d) 4000 masl. Legend: cym_amt = amount of p-cymene (in microlitres); model = floral model used; species = species of flower that the floral model is representing; pf_amt = amount of 2-pentylfuran (in microlitres); pin_amt = amount of α-pinene (in microlitres).

**Table S1.a.** Floral VOCs identified in the first phase of floral VOC identification of the four flower species, viz. *Anemone obtusiloba, Bistorta vivipara, Fragaria nubicol*a and *Lysimachia prolifera*. Floral VOCs were initially identified based only on the mass fragmentation pattern and the retention time (RT) of the compounds. The peaks that could not be identified were labelled as NI (Not Identified), along with their respective m/z (Mass/Charge) values.

| **Flower species** | **m/z** | **RT** | **Floral VOC** |
|---|---|---|---|
| *Anemone obtusiloba* | 193 | 8.27 | NI193 |
| | 133 | 9.43 | NI133 |
| | 67 | 9.62 | cis-3-Hexenyl acetate |
| | 119 | 10.08 | p-cymene |
| | 93 | 10.18 | d-limonene |
| | 93 | 10.79 | trans-b-ocimene |
| | 223 | 10.91 | NI223 |
| | 208 | 11.14 | NI208 |
| | 208 | 11.93 | NI208a |
| | 126 | 12.81 | NI126 |
| | 119 | 14.91 | a-terpineol |
| | 193 | 16.22 | NI193a |
| | 173 | 16.65 | benzaldehyde |
| | 173 | 19.02 | acetophenone |
| | 57 | 19.87 | farnesane |
| | 161 | 20.67 | longifolene |
| | 132 | 30.25 | NI132 |
| | 119 | 30.61 | a-cedrene |
| *Bistorta vivipara* | 91 | 8.09 | Isocumene |
| | 193 | 8.26 | NI193 |
| | 106 | 8.27 | benzaldehyde |
| | 81 | 9.16 | 2-Pentylfuran |
| | 133 | 9.43 | NI133 |
| | 56 | 9.48 | octanal |
| | 68 | 10.18 | d-limonene |
| | 93 | 10.78 | trans-b-ocimene |
| | 223 | 10.92 | NI223 |
| | 208 | 11.14 | NI208a |
| | 208 | 11.95 | NI208 |
| | 126 | 12.82 | NI126 |
| | 152 | 12.98 | NI152 |
| | 193 | 14.56 | NI193a |
| | 193 | 16.26 | NI193b |
| | 208 | 17.36 | NI208b |
| | 105 | 18.66 | NI105 |
| | 173 | 19.02 | acetophenone |

| | | | |
|---|---|---|---|
| | 161 | 20.68 | longifolene |
| | 161 | 22.49 | Germacrene-D |
| | 93 | 29.29 | NI93 |
| | 145 | 29.72 | aromadendrene,dehydro |
| | 149 | 29.94 | NI149 |
| | 58 | 30.17 | NI58 |
| | 119 | 30.61 | a-cedrene |
| *Fragaria nubicola* | 177 | 7.36 | NI177 |
| | 91 | 8.07 | Isocumene |
| | 193 | 8.25 | NI193 |
| | 81 | 9.15 | 2-Pentylfuran |
| | 57 | 9.47 | Octanal |
| | 67 | 9.64 | cis-3-Hexenyl acetate |
| | 93 | 9.65 | d-carene |
| | 119 | 10.07 | m-cymene |
| | 68 | 10.19 | Limonene |
| | 81 | 10.27 | p-cineole |
| | 93 | 10.50 | a-pinene |
| | 223 | 10.92 | NI223 |
| | 83 | 11.61 | NI83 |
| | 71 | 12.94 | farnesane |
| | 152 | 12.97 | 4-pentenylcyclohexane |
| | 69 | 13.21 | NI69 |
| | 133 | 13.52 | dimethylcumene |
| | 193 | 14.56 | NI193a |
| | 193 | 16.25 | NI193b |
| | 131 | 16.49 | NI131 |
| | 173 | 16.65 | benzaldehyde |
| | 208 | 17.36 | NI208 |
| | 105 | 18.66 | Isobutyl benzoate |
| | 173 | 19.01 | acetophenone |
| | 161 | 20.67 | longifolene |
| | 59 | 21.26 | NI59 |
| | 165 | 21.98 | NI165 |
| | 161 | 22.49 | Germacrene D |
| | 173 | 23.77 | benzoic acid |
| | 153 | 24.30 | 2,5-Diisobutyl thiophene |
| | 166 | 24.88 | NI166 |
| | 145 | 29.72 | NI145 |
| | 132 | 30.25 | NI132 |
| | 119 | 30.61 | a-cedrene |
| *Lysimachia prolifera* | 91 | 8.09 | Isocumene |
| | 193 | 8.28 | NI193b |
| | 81 | 9.16 | 2-Pentylfuran |
| | 67 | 9.62 | cis-3-Hexenyl acetate |

| | | |
|---|---|---|
| 119 | 10.07 | m-cymene |
| 93 | 10.18 | Limonene |
| 208 | 11.99 | NI208b |
| 152 | 12.96 | pulegone |
| 193 | 16.22 | NI193 |
| 173 | 16.65 | benzaldehyde |
| 208 | 17.36 | NI208 |
| 173 | 19.07 | acetophenone |
| 71 | 19.27 | NI71 |
| 57 | 19.86 | farnesane |
| 161 | 20.67 | longifolene |
| 59 | 21.26 | NI59a |
| 59 | 23.96 | NI59 |
| 132 | 30.25 | NI132 |

**Table S1.b.** Revised floral Volatile Organic Compounds (VOCs) detected in each of the four flower species with their respective calculated and predicted RRIs (Relative Retention Indices), base peaks, and presence (P)/absence(A) at the elevations where the floral samples were collected. Elevations at which the flower species were not sampled due to low numbers are marked ‘na’.

| **Flower species** | **Floral VOCs** | **Calculated RRI** | **Predicted RRI** | **Base peak (m/z)** | **Presence/Absence at elevations (masl)** | | |
|---|---|---|---|---|---|---|---|
| | | | | | **3000** | **3500** | **4000** |
| *Bistorta vivipara* | Isocumene | 953 | 953 | 91 | P | P | P |
| | 2-Pentylfuran | 992 | 993 | 81 | P | P | P |
| | Limonene | 1028 | 1030 | 93 | P | P | P |
| | trans-b-ocimene | 1049 | 1049 | 119 | A | P | A |
| | Longifolene | 1408 | 1405 | 161 | P | A | P |
| | Germacrene D | 1481 | 1481 | 161 | P | A | A |
| *Anemone obtusiloba* | cis-3-Hexenyl Acetate | 1009 | 1005 | 67 | na | P | P |
| | p-cymene | 1024 | 1025 | 119 | na | P | A |
| | Linomene | 1028 | 1030 | 93 | na | P | P |
| | trans-b-ocimene | 1049 | 1049 | 93 | na | A | P |
| | a-terpineol | 1193 | 1189 | 59 | na | P | A |
| | Longifolene | 1408 | 1405 | 161 | na | P | P |
| *Fragaria nubicola* | Isocumene | 952 | 953 | 91 | P | P | P |
| | 2-Pentylfuran | 992 | 993 | 81 | P | P | P |
| | Octanal | 1003 | 1003 | 56 | P | P | P |
| | cis-3-Hexenyl Acetate | 1009 | 1005 | 67 | P | P | P |
| | d-3-carene | 1010 | 1011 | 93 | P | P | P |
| | m-cymene | 1024 | 1023 | 119 | P | P | P |
| | Limonene | 1028 | 1030 | 93 | P | P | P |
| | p-cineole | 1031 | 1032 | 81 | P | A | A |
| | NI83 | 1077 | - | 83 | P | P | P |
| | 4-pentenylcyclohexane | 1125 | 1124 | 152 | P | P | P |
| | NI69 | 1133 | - | 69 | P | P | P |

| | | | | | | | |
|---|---|---|---|---|---|---|---|
| | isobutyl benzoate | 1331 | 1321 | 105 | A | P | A |
| | Longifolene | 1408 | 1405 | 161 | P | P | P |
| | NI59 | 1432 | - | 59 | P | P | P |
| | NI165 | 1461 | - | 165 | P | A | A |
| | Germacrene D | 1482 | 1481 | 161 | P | A | A |
| | 2,5-Diisobutyl thiophene | 1530 | 1533 | 153 | P | A | A |
| | NI166 | 1542 | - | 166 | P | P | P |
| *Lysimachia prolifera* | Isocumene | 953 | 953 | 91 | P | P | na |
| | 2-Pentylfuran | 992 | 993 | 81 | P | P | na |
| | cis-3-Hexenyl Acetate | 1009 | 1005 | 67 | A | P | na |
| | m-cymene | 1024 | 1023 | 119 | P | P | na |
| | Limonene | 1028 | 1030 | 93 | P | P | na |
| | 4-pentenylcyclohexane | 1124 | 1124 | 95 | P | P | na |
| | Farnesane | 1377 | 1366 | 57 | P | P | na |
| | Longifolene | 1408 | 1405 | 161 | P | P | na |

**Table S2.** Standardised proportions used to paint the floral models

| Floral model | Standardised paint blend using Camlin acrylic paints (KOKUYO CO., LTD.; Japan) |
|---|---|
| AO3500, AO4000 | first white coat then multiple coats of 6 parts Yellow + 1 part White + 1 part Red + 0.25 parts Green + touch of Blue |
| FN3000, FN3500, FN4000 | centre-no white coat-8 parts of Yellow + 1 part Indian red + 0.5 parts Green + 0.5 parts of Blue-one coat; petals painted with one-two coats of White |
| LP3000, LP3500 | first white coat then multiple coats of 3 parts White + 0.5 parts Indian red + 0.5 parts Orange + 0.5 parts Green |

**Table S3.** Details of the blend components/constituent chemicals

| Chemical | IUPAC name | Chemical class | Formula | Number of carbon atoms | Solubility in water | Boiling point (°C) (Sigma-Aldrich) | Melting point (°C) (Sigma-Aldrich) | Density (Sigma Aldrich) | Vapour pressure (mmHg @ 25.00 °C) (goodscentscompany.com) | Approximate Retention Time (min) in the method used | Predicted Relative Retention Index (RRI, HP-5 column, from PheroBase) |
|---|---|---|---|---|---|---|---|---|---|---|---|
| alpha-pinene | (1S,5S)-2,6,6-Trimethylbicyclo[3.1.1]hept-2-ene ((−)-α-Pinene) | terpene | $C_{10}H_{16}$ | 10 | insoluble | 155-156 | -62.5 | 0.858 g/mL at 25 °C (lit.) | 4.75 | 6.9-7 | 982 |
| 2-pentylfuran | 2-pentylfuran | ether | $C_9H_{14}O$ | 9 | slightly | 64-66 | NA | 0.883 g/mL at 20 °C (lit.), 0.886 g/mL at 25 °C (lit.) | 2.022 | 9.7-9.9 | 1001 |
| 3-carene | 3,7,7-trimethylbicyclo[4.1.0]hept-3-ene | terpene | $C_{10}H_{16}$ | 10 | insoluble | 168-169 | < 25 °C | 0.857 g/mL at 25 °C (lit.) | 1.862 | 10.8-10.9 | 1011 |
| cis-3-hexenyl acetate | [(Z)-hex-3-enyl] acetate | acetate ester | $C_8H_{14}O_2$ | 8 | NA | 165-167 | NA | 0.897 g/mL at 25 °C (lit.) | 1.219 | 11.3-11.4 | 1007 |
| p-cymene | 1-methyl-4-propan-2-ylbenzene | monoterpene | $C_{10}H_{14}$ | 10 | insoluble | 176-178 | -68 °C | 0.86 g/mL at 25 °C (lit.) | 1.46 | 11.2-11.4 | 1027 |

| | | | | | | | | | | | |
|---|---|---|---|---|---|---|---|---|---|---|---|
| D-limonene | (4R)-1-methyl-4-prop-1-en-2-ylcyclohexene | terpene | $C_{10}H_{16}$ | 10 | insoluble | 176-177 | -74 °C | 0.842 g/mL at 20 °C (lit.) | 0.198 | 11.4-11.5 | 1031 |
| alpha-cedrene | (1S,2R,5S,7S)-2,6,6,8-tetramethyltricyclo[5.3.1.01,5]undec-8-ene | terpene | $C_{15}H_{24}$ | 15 | NA | 261-262 | NA | 0.932 g/mL at 20 °C (lit.) | 0.018 | 33.1-33.3 | 1410 |

**Table S4.** Method for reconstitution of blends: Out of the blend components, the compound with the highest percentage was considered to have a value of 100%. Accordingly, the percentages of the other blend components with respect to this component were calculated. Thus, the following table of the expected relative ratios of the blend components were obtained.

| **Blend name** | **Chemical constituent** | **Minimum percentage emission** | **Maximum percentage emission** | **Minimum Relative Ratio** | **Maximum Relative Ratio** | **Amount to be added (µl) for a $10^{-1}$ concentration** |
|---|---|---|---|---|---|---|
| AO3500 | 3-carene | 0 | 0.19 | 0.00 | 7.42 | 1 |
| | cis-3-hexenyl acetate | 0.52 | 2.56 | 20.31 | 100.00 | 15 |
| | p-cymene | 0.07 | 0.17 | 2.73 | 6.64 | 0.05 |
| | D-limonene | 0.09 | 0.53 | 3.52 | 20.70 | 0.66 |
| | alpha-cedrene | 0 | 0.74 | 0.00 | 28.91 | 500 |
| AO4000 | 3-carene | 0 | 0.149 | 0.00 | 3.04 | 0.1 |
| | cis-3-hexenyl acetate | 0.29 | 3.75 | 5.92 | 76.53 | 7.5 |
| | p-cymene | 0 | 0.22 | 0.00 | 4.49 | 0.1 |
| | D-limonene | 0.51 | 2.35 | 10.41 | 47.96 | 0.5 |
| | alpha-cedrene | 0 | 4.9 | 0.00 | 100.00 | 10 |
| FN3000 | alpha-pinene | 0 | 0 | 0.00 | 0.00 | 0 |
| | 2-pentylfuran | 0.45 | 1.52 | 7.04 | 23.79 | 23 |

| | | | | | | |
|---|---|---|---|---|---|---|
| | 3-carene | 0.15 | 1.07 | 2.35 | 16.74 | 16 |
| | cis-3-hexenyl acetate | 0 | 0.25 | 0.00 | 3.91 | 10 |
| | p-cymene | 0.27 | 1.21 | 4.23 | 18.94 | 18 |
| | D-limonene | 1.27 | 6.39 | 19.87 | 100.00 | 100 |
| | alpha-cedrene | 0 | 3.06 | 0.00 | 47.89 | 50 |
| FN3500 | alpha-pinene | 0.03 | 0.09 | 1.65 | 4.95 | 0.5 |
| | 2-pentylfuran | 0 | 0.28 | 0.00 | 15.38 | 1.5 |
| | 3-carene | 0 | 0 | 0.00 | 0.00 | 0 |
| | cis-3-hexenyl acetate | 0.8 | 1.82 | 43.96 | 100.00 | 100 |
| | p-cymene | 0.05 | 0.15 | 2.75 | 8.24 | 0.8 |
| | D-limonene | 0.03 | 0.41 | 1.65 | 22.53 | 2.2 |
| | alpha-cedrene | 0 | 0.64 | 0.00 | 35.16 | 35 |
| FN4000 | alpha-pinene | 0 | 0.01 | 0.00 | 0.22 | 0.5 |
| | 2-pentylfuran | 0 | 0.29 | 0.00 | 6.36 | 9 |
| | 3-carene | 0 | 0.07 | 0.00 | 1.54 | 5 |

| | | | | | | |
|---|---|---|---|---|---|---|
| | cis-3-hexenyl acetate | 0.48 | 4.56 | 10.53 | 100.00 | 40 |
| | p-cymene | 0.03 | 0.13 | 0.66 | 2.85 | 6 |
| | D-limonene | 0.12 | 0.44 | 2.63 | 9.65 | 4 |
| | alpha-cedrene | 0 | 0.9 | 0.00 | 19.74 | 170 |
| LP3000 | 2-pentylfuran | 0.03 | 0.85 | 2.21 | 62.50 | 0.75 |
| | cis-3-hexenyl acetate | 0 | 0.91 | 0.00 | 66.91 | 7.5 |
| | p-cymene | 0 | 0.55 | 0.00 | 40.44 | 0.5 |
| | D-limonene | 0.42 | 1.36 | 30.88 | 100.00 | 10 |
| LP3500 | 2-pentylfuran | 0.01 | 0.15 | 0.45 | 6.70 | 1 |
| | cis-3-hexenyl acetate | 1.04 | 2.24 | 46.43 | 100.00 | 1000 |
| | p-cymene | 0.03 | 0.07 | 1.34 | 3.13 | 0.5 |
| | D-limonene | 0.06 | 0.3 | 2.68 | 13.39 | 3 |

**Table S5.** Summary table of results of chi-square tests of independence of cafeteria assays

| S. no. | Comparison | Chi-square value | p-value | Sample size | Degrees of freedom |
|---|---|---|---|---|---|
| (i) | Sikkim_May_VisualCues_3000masl_ vs. null model (no preference) | 54.379 | <0.0001 | 166 | 6 |
| (ii) | Sikkim_May_VisualCues_3500masl_ vs. null model (no preference) | 7.4 | 0.2854 | 91 | 6 |
| (iii) | Sikkim_May_VisualCues_4000masl_ vs. null model (no preference) | 61.348 | <0.0001 | 161 | 6 |
| (iv) | Sikkim_May_OlfactoryandVisualCues_3000masl_ vs. null model (no preference) | 33.326 | <0.0001 | 105 | 6 |
| (v) | Sikkim_May_OlfactoryandVisualCues_3500masl_ vs. null model (no preference) | 35.663 | <0.0001 | 148 | 6 |
| (vi) | Sikkim_May_OlfactoryandVisualCues_4000masl_ vs. null model (no preference) | 73.198 | <0.0001 | 182 | 6 |
| (vii) | Sikkim_Sep_VisualCues_3000masl_ vs. null model (no preference) | 36.909 | <0.0001 | 203 | 6 |
| (viii) | Sikkim_Sep_VisualCues_3500masl_ vs. null model (no preference) | 32.407 | <0.0001 | 123 | 6 |
| (ix) | Sikkim_Sep_VisualCues_4000masl_ vs. null model (no preference) | 41.768 | <0.0001 | 149 | 6 |
| (x) | Sikkim_Sep_OlfactoryandVisualCues_3000masl_ vs. null model (no preference) | 28.221 | <0.0001 | 139 | 6 |
| (xi) | Sikkim_Sep_OlfactoryandVisualCues_3500masl_ vs. null model (no preference) | 31.908 | <0.0001 | 147 | 6 |
| (xii) | Sikkim_Sep_OlfactoryandVisualCues_4000masl_ vs. null model (no preference) | 25.091 | 0.0003 | 97 | 6 |
| (xiii) | Bengaluru_VisualCues_ vs. null model (no preference) | 17.958 | 0.0063 | 475 | 6 |
| (xiv) | Bengaluru_OlfactoryandVisualCues_ vs. null model (no preference) | 6.628 | 0.3566 | 316 | 6 |
| (xv) | Sikkim_May_OlfactoryandVisualCues_3000masl_ vs. Sikkim_May_OlfactoryandVisualCues_3500masl_ | 16.874 | < 0.0315 | 253 | 13 |

| | | | | | |
|---|---|---|---|---|---|
| (xvi) | Sikkim_May_OlfactoryandVisualCues_3500masl_<br>vs.<br>Sikkim_May_OlfactoryandVisualCues_4000masl_ | 50.305 | <0.0001 | 330 | 13 |
| (xvii) | Sikkim_May_OlfactoryandVisualCues_3000masl_<br>vs.<br>Sikkim_May_OlfactoryandVisualCues_4000masl_ | 32.352 | <0.0001 | 287 | 13 |
| (xviii) | Sikkim_Sep_OlfactoryandVisualCues_3000masl_<br>vs.<br>Sikkim_Sep_OlfactoryandVisualCues_3500masl_ | 5.655 | 0.6859 | 286 | 13 |
| (xix) | Sikkim_Sep_OlfactoryandVisualCues_3500masl_<br>vs.<br>Sikkim_Sep_OlfactoryandVisualCues_4000masl_ | 21.279 | 0.0064 | 244 | 13 |
| (xx) | Sikkim_Sep_OlfactoryandVisualCues_3000masl_<br>vs.<br>Sikkim_Sep_OlfactoryandVisualCues_4000masl_ | 36.759 | <0.0001 | 236 | 13 |
| (xxi) | Sikkim_May_OlfactoryandVisualCues_3000masl_<br>vs.<br>Sikkim_Sep_OlfactoryandVisualCues_3000masl_ | 18.663 | 0.0048 | 244 | 13 |
| (xxii) | Sikkim_May_OlfactoryandVisualCues_3500masl_<br>vs.<br>Sikkim_Sep_OlfactoryandVisualCues_3500masl_ | 23.594 | 0.0006 | 295 | 13 |
| (xxiii) | Sikkim_May_OlfactoryandVisualCues_4000masl_<br>vs.<br>Sikkim_Sep_OlfactoryandVisualCues_4000masl_ | 11.039 | 0.0872 | 279 | 13 |
| (xxiv) | Sikkim_May_VisualCues_3000masl_<br>vs.<br>Sikkim_May_OlfactoryandVisualCues_3000masl_ | 9.052 | 0.3379 | 166 | 13 |
| (xxv) | Sikkim_May_VisualCues_3500masl_<br>vs.<br>Sikkim_May_OlfactoryandVisualCues_3500masl_ | 24.562 | 0.0018 | 91 | 13 |
| (xxvi) | Sikkim_May_VisualCues_4000masl_<br>vs.<br>Sikkim_May_OlfactoryandVisualCues_4000masl_ | 15.068 | 0.0578 | 161 | 13 |

185

**Table S6.** Summary of the chemicals collected from six types of blooming flowers during behavioural assays in Bengaluru using Polydimethylsiloxane (PDMS) tubes and identified using Thermal Desorption Unit-Gas Chromatography-Mass Spectrometry (TDU-GCMS). The relative retention index (RRI) was calculated against a hydrocarbon standard run using the same method as the floral VOC analyses. The first and second highest m/z peak values of the detected chemicals are also listed.

| **Sample ID** | **Flower type** | **Retention time (minutes)** | **Calculated RRI** | **m/z highest peak** | **m/z second highest peak** |
|---|---|---|---|---|---|
| YF1 | Yellow flower 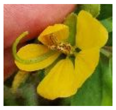 | 4.7 | 900 | 77 | 45 |
| | | 6.25 | 970.4545455 | 207 | 96 |
| | | 6.9 | 996.9162996 | 207 | 208 |
| | | 9 | 1076.315789 | 207 | 75 |
| | | 10.56 | 1034.782609 | 133 | 151 |
| | | 12 | 1186.813187 | 106 | 105 |
| | | 15.12 | 1303.137255 | 281 | 282 |
| | | 20.5 | 1620.434783 | 207 | 208 |
| | | 21.46 | 1662.173913 | 41 | 57 |
| | | 25.5 | 1741.037736 | 73 | 355 |
| | | 33 | 2711.174785 | 142 | 141 |
| WF1 | White flower 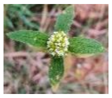 | 4.73 | 901.3636364 | 77 | 45 |
| | | 6.23 | 969.5454545 | 207 | 96 |
| | | 10.52 | 1033.333333 | 133 | 151 |
| | | 12.67 | 1211.567164 | 194 | 209 |
| | | 15.11 | 1302.745098 | 281 | 282 |
| | | 25.51 | 1741.509434 | 73 | 355 |
| | | 34.55 | 2755.587393 | 73 | 341 |
| LF1 | Lantana flower 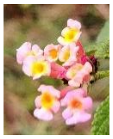 | 4.51 | 891.3636364 | 77 | 45 |
| | | 6.28 | 971.8181818 | 207 | 96 |
| | | 10.32 | 1026.086957 | 133 | 151 |
| | | 11.14 | 1055.797101 | 179 | 153 |
| | | 14.77 | 1389.925373 | 146 | 148 |
| | | 15.13 | 1303.529412 | 281 | 282 |
| | | 20.56 | 1623.043478 | 207 | 96 |
| | | 21.47 | 1662.608696 | 41 | 57 |
| | | 25.51 | 1741.509434 | 73 | 355 |
| | | 31.03 | 2060.992908 | 146 | 148 |
| PFC1 | Purple Flower (Conventional) 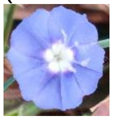 | 5.02 | 914.5454545 | 77 | 45 |
| | | 6.24 | 970 | 207 | 96 |
| | | 10.67 | 1038.768116 | 133 | 151 |
| | | 15.12 | 1303.137255 | 281 | 282 |
| | | 20.6 | 1624.782609 | 207 | 96 |
| | | 21.46 | 1662.173913 | 41 | 57 |
| | | 25.51 | 1741.509434 | 73 | 355 |
| | | 34.55 | 2355.587393 | 73 | 341 |
| YFN1 | Yellow Flower (New) 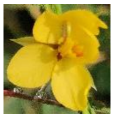 | 4.7 | 900 | 77 | 45 |
| | | 6.25 | 970.4545455 | 207 | 96 |
| | | 10.46 | 1031.15942 | 133 | 151 |
| | | 15.12 | 1303.137255 | 281 | 282 |
| | | 20.59 | 1624.347826 | 207 | 96 |

| | | | | | |
|---|---|---|---|---|---|
| | | 25.5 | 1741.037736 | 73 | 355 |
| | | 30.8 | 2044.366197 | 281 | 282 |
| | | 31.02 | 2006.666667 | 146 | 148 |
| | | 34.55 | 2355.587393 | 73 | 341 |
| PFB1 | Purple Flower (Bristly) 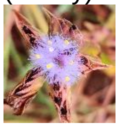 | 906.3636364 | 906.3636364 | 77 | 45 |
| | | 970.9090909 | 970.9090909 | 207 | 96 |
| | | 1035.144928 | 1035.144928 | 133 | 151 |
| | | 1303.529412 | 1303.529412 | 281 | 282 |
| | | 1622.173913 | 1622.173913 | 207 | 96 |
| | | 1662.173913 | 1662.173913 | 41 | 57 |
| | | 1741.509434 | 1741.509434 | 73 | 355 |
| | | 2045.774648 | 2045.774648 | 281 | 282 |
| | | 2006.666667 | 2006.666667 | 146 | 148 |
| | | 2355.587393 | 2355.587393 | 73 | 341 |

**Table S7: Summary table of results of generalised linear models (GLMs)**

| **Response variable** | **Predictor variable** | **Estimate** | **p-value** | **AIC** | **Null deviance** | **Residual deviance** | **Pseudo-$R^2$** |
|---|---|---|---|---|---|---|---|
| Visits to floral model across elevations | Amount of α-pinene | 1.62186 | 6.99E-14 | 791.26 | 496.81 | 442.95 | 0.108411666 |
| Visits to floral model across elevations | Amount of 2-pentylfuran | 0.04014 | 7.98E-13 | 799.25 | 496.81 | 450.94 | 0.092329059 |
| Visits to floral model across elevations | Amount of 3-carene | 0.0502 | 4.68E-10 | 810.49 | 496.81 | 462.18 | 0.069704716 |
| Visits to floral model across elevations | Amount of cis-3-hexenyl acetate | -0.0001795 | 0.288 | 843.94 | 496.81 | 495.63 | 0.002375153 |
| Visits to floral model across elevations | Amount of p-cymene | 0.048707 | 9.14E-12 | 803.76 | 496.81 | 455.45 | 0.083251142 |
| Visits to floral model | Amount of D-limonene | 0.006899 | 1.10E-07 | 820.13 | 496.81 | 471.82 | 0.05030092 |

| | | | | | | | |
|---|---|---|---|---|---|---|---|
| across elevations | | | | | | | |
| Visits to floral model across elevations | Amount of α-cedrene | -0.0013077 | 0.000822 | 832.23 | 496.81 | 483.92 | 0.025945532 |
| Visits to floral model at 3000 masl | Amount of α-pinene | 1.88322 | 2.86E-05 | 186.41 | 114.238 | 97.277 | 0.148470737 |
| Visits to floral model at 3000 masl | Amount of 2-pentylfuran | 0.03192 | 0.00802 | 197 | 114.24 | 107.87 | 0.055759804 |
| Visits to floral model at 3000 masl | Amount of 3-carene | 0.03677 | 0.0359 | 199.36 | 114.24 | 110.23 | 0.035101541 |
| Visits to floral model at 3000 masl | Amount of cis-3-hexenyl acetate | -0.0002585 | 0.47498 | 202.83 | 114.24 | 113.69 | 0.004814426 |
| Visits to floral model at 3000 masl | Amount of p-cymene | 0.03777 | 0.0144 | 197.96 | 114.24 | 108.82 | 0.047443978 |

| | | | | | | | |
|---|---|---|---|---|---|---|---|
| Visits to floral model at 3000 masl | Amount of D-limonene | 0.004708 | 0.1001 | 200.9 | 114.24 | 111.77 | 0.021621148 |
| Visits to floral model at 3000 masl | Amount of α-cedrene | -0.0015926 | 0.061521 | 199.2 | 114.24 | 110.07 | 0.036502101 |
| Visits to floral model at 3500 masl | Amount of α-pinene | 1.14669 | 0.00363 | 263.98 | 134.94 | 126.85 | 0.059952572 |
| Visits to floral model at 3500 masl | Amount of 2-pentylfuran | 0.047082 | 1.72E-06 | 251.74 | 134.94 | 114.6 | 0.150733659 |
| Visits to floral model at 3500 masl | Amount of 3-carene | 0.06265 | 7.43E-06 | 254.38 | 134.94 | 117.24 | 0.131169409 |
| Visits to floral model at 3500 masl | Amount of cis-3-hexenyl acetate | -0.0005115 | 0.13693 | 269.54 | 134.94 | 132.41 | 0.018749074 |
| Visits to floral model | Amount of p-cymene | 0.05832 | 2.95E-06 | 252.81 | 134.94 | 115.68 | 0.142730102 |

| | | | | | | | |
|---|---|---|---|---|---|---|---|
| at 3500 masl | | | | | | | |
| Visits to floral model at 3500 masl | Amount of D-limonene | 0.008864 | 7.18E-05 | 258.36 | 134.94 | 121.23 | 0.101600711 |
| Visits to floral model at 3500 masl | Amount of α-cedrene | -0.0009424 | 0.15496 | 269.84 | 134.94 | 132.7 | 0.01659997 |
| Visits to floral model at 4000 masl | Amount of α-pinene | 1.8071 | 1.51E-08 | 322.99 | 223.9 | 192.95 | 0.138231353 |
| Visits to floral model at 4000 masl | Amount of 2-pentylfuran | 0.03935 | 2.10E-06 | 333.8 | 223.9 | 203.76 | 0.089950871 |
| Visits to floral model at 4000 masl | Amount of 3-carene | 0.04794 | 6.26E-05 | 339.6 | 223.9 | 209.56 | 0.064046449 |
| Visits to floral model at 4000 masl | Amount of cis-3-hexenyl acetate | 4.00E-05 | 0.863 | 353.92 | 223.9 | 223.87 | 0.000133988 |

| | | | | | | | |
|---|---|---|---|---|---|---|---|
| Visits to floral model at 4000 masl | Amount of p-cymene | 0.04733 | 7.83E-06 | 336.15 | 223.9 | 206.11 | 0.079455114 |
| Visits to floral model at 4000 masl | Amount of D-limonene | 0.006549 | 0.000711 | 343.74 | 223.9 | 213.7 | 0.045556052 |
| Visits to floral model at 4000 masl | Amount of α-cedrene | -0.001435 | 0.0148 | 347 | 223.9 | 216.96 | 0.03099598 |

**Dataset S1:** Dataset of floral Volatile Organic Compounds (VOCs) detected in each of the four species, viz., *Bistorta vivipara*, *Anemone obtusiloba*, *Fragaria nubicola* and *Lysimachia prolifera*, and their respective Calculated and Predicted RRIs (Relative Retention Index), base peaks, and presence (P)/absence(A) at the elevations where the floral samples were collected. Elevations from where samples were not collected are marked 'na' (not applicable). Non-identified VOCs are marked as NIs (not identified) along with their respective mass/charge in the GC-MS output.

**Dataset S2:** Dataset of wild pollinator visitation counts received by all artificial floral models (1) in cafeteria assays performed at 3000 masl, 3500 masl and 4000 masl in Sikkim in flowering seasons of May and September, and novel plant-pollinator communities in Bengaluru, (2) in subtractive assays at 3000 masl, 3500 masl and 4000 masl in Sikkim.
